# Do persistent rare species experience stronger negative frequency dependence than common species?

**DOI:** 10.1101/040360

**Authors:** Glenda Yenni, Peter B. Adler, S. K. Morgan Ernest

**Affiliations:** Department of Wildlife Ecology and Conservation, University of Florida, Gainesville, FL 32611; Department of Wildland Resources, Utah State University, Logan, UT, 84322

**Keywords:** rare species, coexistence, frequency dependence

## Abstract

Understanding why so many species are rare yet persistent remains a significant challenge for both theoretical and empirical ecologists. Yenni, Adler, and Ernest (2012) proposed that strong negative frequency dependence causes species to be rare while simultaneously buffering them against extinction. This hypothesis predicts that, on average, rare species should experience stronger negative frequency dependence than common species. However, it is unknown if ecological communities generally show this theoretical pattern, or if rarity is primarily determined by other processes that overwhelm the effects of strong negative frequency dependence. We discuss the implications of this mechanism for natural communities, and develop a method to test for a non-random relationship between negative frequency dependence and relative abundance, using species abundance data from 90 communities across a broad range of environments and taxonomic groups. To account for biases introduced by measurement error, we compared the observed correlation between species relative abundance and the strength of frequency dependence against expectations from a randomization procedure. In approximately half of the analyzed communities, rare species showed disproportionately strong negative frequency dependence compared to common species. Specifically, we found a pattern of increasingly strong negative frequency dependence with decreasing relative abundance. Our results suggest that strong negative frequency dependence is a signature of both rarity and persistence for many species in many communities.

## Introduction

Rare species are ubiquitous in nature, but understanding the factors limiting their abundance and the mechanisms allowing them to persist has proven challenging. Reproducing realistic local-scale community structures with numerous rare species remains difficult using simple theoretical or statistical models of species interactions (Tilman, 2004; Angert *et al.*, 2009; Clark *et al.*, 2010). One reason these models fail to predict realistic numbers of rare species is that many rare species are simply ephemeral, and their occasional presence is likely explained by metapopulation dynamics. Thus the number of these species in a local community can only be explained when regional-scale processes are considered along with local-scale processes. However, not all rare species are ephemeral. Some species manage to persist at low abundance within a community for long periods of time without succumbing to stochastic extinction events. Since stochasticity is pervasive in ecological systems, species with low population sizes need mechanisms that buffer their population dynamics from extinction in order to persist (Tilman, 2004; Kobe & Vriesendorp, 2011). It is this subset of rare species, the persistent rare species, which remain a paradox for coexistence. Despite the importance of understanding how these rare species are able to persist, how to determine a rare species’ role in a community as persistent or ephemeral, how prevalent persistent rare species are, and what allows their persistence all remain controversial (Preston, 1948; Main, 1982; Kunin & Gaston, 1997; Magurran & Henderson, 2003; Comita *et al.*, 2010). Here we provide empirical evidence for a recently proposed mechanism that, in theory, can buffer small populations against extinction: asymmetric negative frequency dependence (Yenni *et al.*, 2012).

## The role of Negative Frequency Dependence in rarity and persistence

Like density dependence, frequency dependence is a relationship between the population growth rate and the frequency of a species in a community. However, because it is based on relative frequency (i.e. abundance relative to other community members) rather than absolute abundance, frequency dependence arises through both intra and inter-specific interactions (Adler *et al.*, 2007). Negative frequency dependence occurs when a species’ population growth rate is more strongly limited by the density of conspecifics than by the density of heterospecifics (Fig 1A). When negative frequency dependence is strong, relatively small increases in a species’ density, and in turn its frequency in the community, will result in large declines in population growth rate (Fig 1B). Holding the maximum population growth rate constant, species with stronger negative frequency dependence must have lower relative abundance than species with weak negative frequency dependence. In other words, for species experiencing strong negative frequency dependence, intraspecific competition is the primary cause of rarity (Appendix S2 in Supporting Information).

**Figure 1:**
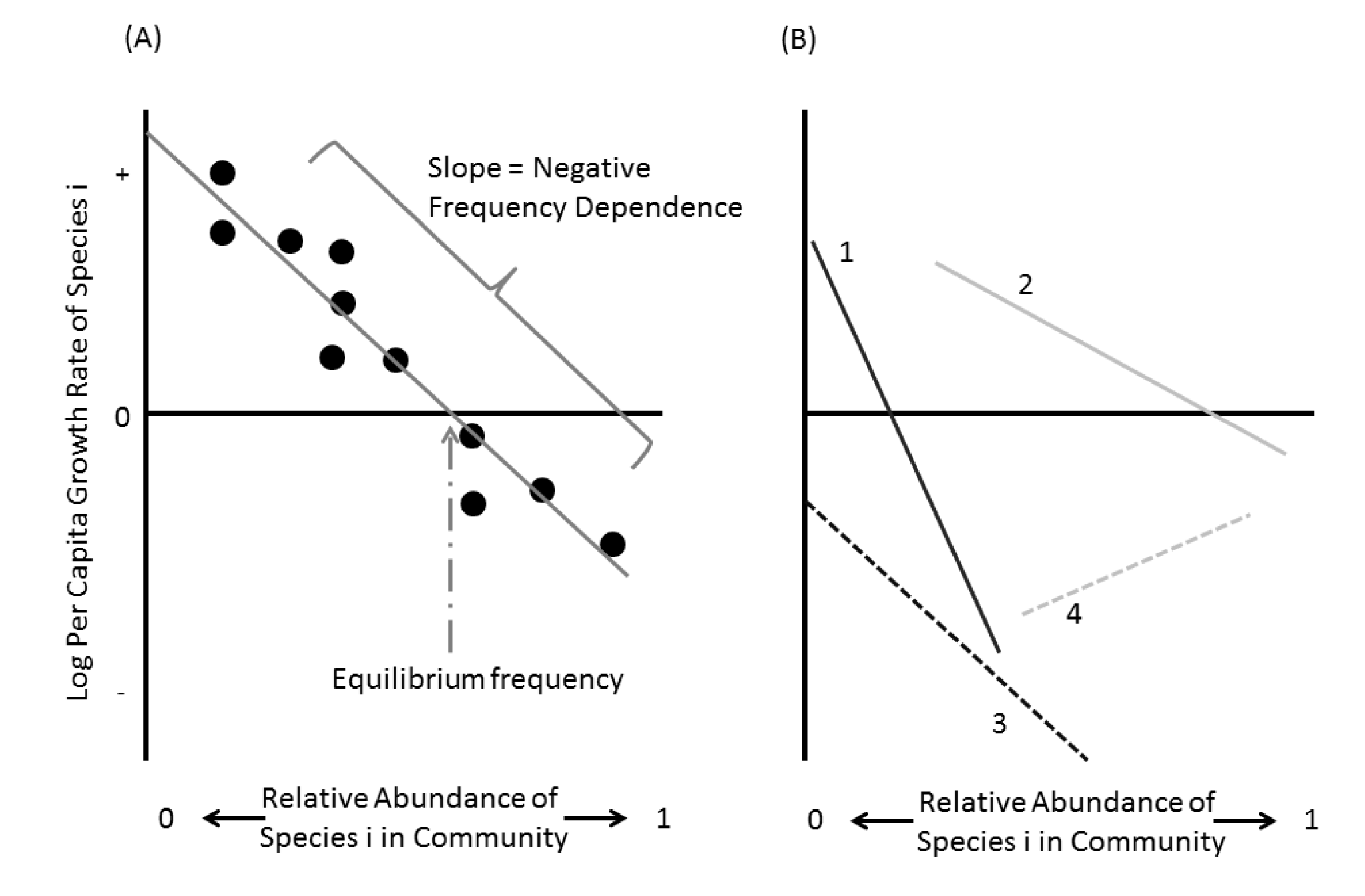
A) Negative frequency dependence and equilibrium frequency can be calculated from time-series community data. Each data point represents the relative abundance of species *i* at time *t* in its community (x-axis) and its log per capita population growth rate from time *t* to time *t+1* (y-axis). The slope of this line is the species’ negative frequency dependence (*NFD* = -β1 in the linear model). Equilibrium frequency – the expected relative abundance of species i in the community when its population growth is at equilibrium - is the fitted x-intercept of the relationship. The fitted y-intercept is the species’ maximal growth rate. B) The x-intercept and slope of the relationship indicate important biological information about each species. Steep slopes indicate stronger negative frequency dependence, and thus higher sensitivity of population growth to changes in conspecifics than to changes in heterospecifics. In a linear relationship, high maximal growth rates, which buffer species from extinction, are achieved through high equilibrium frequencies (common species, solid grey line), or strong negative frequency dependence (rare species, solid black line). X-intercepts < 0 indicate species that cannot maintain a stable presence within the community at their population equilibrium. Species exhibiting positive frequency dependence will often have difficulty invading when rare. These latter two scenarios indicate species that are likely not stably coexisting within the current community.

In contrast, studies of competition often assume that rare species are primarily suppressed by interspecific competition (Chesson, 2000; Tilman, 2004). But if interspecific competition is the cause of rarity, how can we explain the long-term persistence of many rare species? Why don’t the superior competitors exclude rare species altogether?Our alternative explanation, that rare species are strongly frequency-dependent, also explains persistence, because strong population regulation buffers species against extinction. It is initially counter-intuitive that a process causing rarity could also promote persistence. However, when a species with strong negative frequency dependence drops below its equilibrium frequency, it will experience a large increase in population growth rate (Fig. 1). Because species with strong negative frequency dependence are more sensitive to increases in their own population numbers, they experience relatively weak suppression by dominant community members; they are effectively released from competitive effects as their densities approach zero. Thus, strong negative frequency dependence can simultaneously limit species’ average abundances while buffering them against extinction if they fall to low density (Yenni *et al.*, 2012).

Strong negative frequency dependence should benefit rare species more than common species (Yenni *et al.*, 2012). Common species are less vulnerable to stochastic extinction due to their higher abundances. While common species might experience strong or weak negative frequency dependence, rare species lacking strong negative frequency dependence would be quickly lost from a local community due to stochastic extinction. In addition, if a rare and common species experience the same degree of negative frequency dependence (equivalent slopes in Fig 1), then the rare species must have a lower maximal growth rate. Holding the equilibrium abundances constant, for the rare species to experience the same buffering against stochastic extinction (i.e. similar maximal growth rates), it must have stronger negative frequency dependence. Therefore, we expect a relationship between relative abundance (frequency) and the strength of negative frequency dependence: species with lower relative abundances should experience stronger negative frequency dependence.

## The Asymmetric Negative Frequency Dependence Pattern

While simulations suggest that strong negative frequency dependence should be a common feature of persistent rare species (Yenni *et al.*, 2012), few empirical studies have assessed this mechanism. Strong negative density or frequency dependence in some rare species has been documented (Harpole & Suding, 2007; Adler *et al.*, 2010; Comita *et al.*, 2010; Mangan *et al.*, 2010; Johnson *et al.*, 2012; Xu *et al.*, 2015) but these few studies exclusively focused on select plant communities. If this is a general pattern across taxa and ecosystems, then this mechanism could be of general importance to the coexistence and persistence of rare species. However, this limited number of studies does not tell us how general this pattern is across a variety of natural communities. Testing the asymmetric negative frequency dependence hypothesis is challenging because there is no a priori cut-off value for what constitutes strong enough negative frequency dependence to buffer species against stochastic extinction. However, we can examine whether rarer species have stronger negative frequency dependence than more common species, which is equivalent to testing whether rare species are more sensitive to intra‐ than interspecific competition when compared to common species. If strong negative frequency dependence is more common among rare species, it would suggest 1) that rare species are not rare because they are suppressed by superior competitors, but rather because they are more strongly self-limiting, and 2) that rare species that do not experience a large release from competition at low density, as implied by strong negative frequency dependence, often do not persist.

## Testing for Asymmetric Negative Frequency Dependence

To explore the relationship between rarity and negative frequency dependence in ecological communities, we decided to assess whether there is a general relationship between the rarity of a species and the strength of negative frequency dependence by examining a large compilation of publicly available time-series on the abundance of all species within one trophic guild. Using this approach to evaluate the relationship between abundance and rarity requires 1) identification of appropriate data, 2) estimating equilibrium frequency and negative frequency dependence for each species in each community, 3) identifying the persistent community, 4) estimating a community-level pattern of negative frequency dependence, and 5) accounting for known and unknown biases in the pattern of interest before making a final conclusion about the negative frequency dependence-rarity relationship in each community. The methods we developed to deal with these issues have allowed us to use commonly available data to begin to explore the relationship between negative frequency dependence and rarity. We hope our results will spur future research and collection of even more useful data.

## Identifying Suitable Data for Community Negative Frequency Dependence Patterns

Many of the coexistence mechanisms that generate negative frequency dependence can operate at very small spatial scales (Chesson, 2000). This is also true in the framework defined in Yenni et al. (2012). Therefore, to assess whether persistent rare species have stronger negative frequency dependence than common species, we required communities where the data was collected in a manner consistent with this local-scale concept. We restricted communities to all species within a study belonging to the same trophic level and occurring within a single habitat type as defined by the data collector.If the data collector indicated that the study included multiple sites of different habitat types or multiple trophic levels these data were treated as different communities and analyzed separately. If a study collected data from multiple locations of the same habitat type, we examined those separately. However, to keep a single study area from dominating the results because it had multiple sites, we treated multiple sites in a single study area as habitat replicates and only one covariance was estimated from the entire resulting site data.Finally, to calculate frequency dependence, we used time series data because studies measuring frequency dependence for all species in a community in an experimental manner are rare. All of these data decisions were made to ensure we were only analyzing populations that were likely to be directly interacting and to provide data for a large number and diversity of taxa and ecosystems. Using these criteria, we found suitable data for 90 different communities containing collectively 703 species (Appendix S1 in Supplementary Information). These communities represent six major taxonomic groups (birds, fish, herpetofauna, invertebrates, mammals, and plants), 5 continents, and 3 trophic levels, thus providing a general exploration of whether persistent rare species commonly exhibit stronger negative frequency dependence, and thus, stronger population buffering, than common species.

## Calculating Negative Frequency Dependence and Equilibrium Frequency

Community time series can be used to estimate equilibrium frequency – the relative abundance when a species’ population is in steady state – and the strength of negative frequency dependence for all species that met our criteria for persistence within each community (Fig 1A). We estimated a species’ negative frequency dependence as the linear relationship between its relative abundance (frequency) in a community and its annual per capita population growth rate. For each community, relative abundance in each year *t* for each species *s* was calculated as x*t,s* = N*t,s* / Σ_s_=1:S N*t,s* (where N is a species’ absolute abundance). Log per capita population growth rates in each year *t* for each species *s* were calculated as y*t,s* = *log*(N*t*+*1,s*/N*t,s*). The relationship between these population parameters was described as **Y***s* = β_o,s_ + β_1,s_ **X***s* + **ɛ**_*s*_ for each species *s*. Equilibrium frequency (*f*), a species’ expected relative abundance in the community, is estimated as the x-intercept of this linear relationship, *f* = -β_o,s_/β_1,s_. Frequency dependence (*FD*) is estimated as the slope of the linear relationship, *FD* = β_1,s_.

## Ephemeral vs. Persistent Species

As discussed in the introduction, asymmetric negative frequency dependence is a mechanism for maintaining small populations near their steady state despite stochasticity, and thus only applies to stably coexisting common and rare species. Because other mechanisms are expected to dominate the population dynamics of invading, declining, or transient species, we used three criteria to identify and exclude species with population characteristics that would make them likely to be ephemeral and not stably coexisting (Supplementary Figure S15): 1) too few occurrences to estimate parameters (i.e. few or no adjacent non-zero abundances, the inclusion of this type of species in a dataset is typically an arbitrary decision by the data collectors, and was highly variable between datasets), which indicates a species is likely transient, 2) negative equilibrium frequency (Fig 1B), which indicates the species cannot persist in the community (7.7% of species, (Chesson, 2000; Adler *et al.*, 2007)), and 3) positive frequency dependence (Fig 1B), which is related to a negative invasion growth rate and likely results in either the extinction of that species or an ever increasing invader (11% of species, (Chesson, 2000; Adler *et al.*, 2007)). Because equilibrium frequency is estimated as the x-intercept of the fitted relationship between log per capita growth rate and relative abundance, it is possible for this value to be < 0 or > 1 (Fig 1B). Besides the fact that these species are obviously ephemeral and thus irrelevant to our analysis, it would make the relationship between equilibrium frequency and frequency dependence nonsensical to include negative equilibrium frequencies or positive frequency dependences.

All other species are retained as persistent community members. Our criteria are only a weak filter for removing ephemeral species. Some of the species retained in the analysis are likely also ephemeral, but we abstained from further filtering to avoid possibly biasing the results. We examined how this method of identifying persistent species relates to actual persistence at a site by calculating the percent of years in which a species has non-zero abundance and comparing these values between ephemeral and persistent species. Our approach identified a group of more persistent species that were on average present over a higher fraction of the time series (Supplementary Information Figure S14).

## Quantifying Asymmetric Negative Frequency Dependence at the Community Level

We currently have no way to objectively characterize a species as experiencing weak or strong negative frequency dependence. We can, however, estimate the relationship between relative abundance and negative frequency dependence across all persistent species in the community. To quantify the relationship between rarity and negative frequency dependence within a community, we calculated the covariance between equilibrium frequency and strength of negative frequency dependence (Fig 2B). The covariance calculation uses every species in a community as a single data point in the calculation (Fig 2B): *cov*(*log*(***f***), *log*(***FD***)).Negative covariances indicate communities in which rare species are experiencing stronger negative frequency dependence than their dominant counterparts (note that the expectation for *cov*(*log*(***f***), *log*(***FD***)) is already negative, we discuss how we addressed this in the following section). Log-log relationships were most appropriate for calculating these covariances because most communities contain many species with very low equilibrium frequencies and only a few dominants, and because the observed negative frequency dependence of the few dominants was typically at least an order of magnitude lower than the remainder of the community.

**Figure 2:**
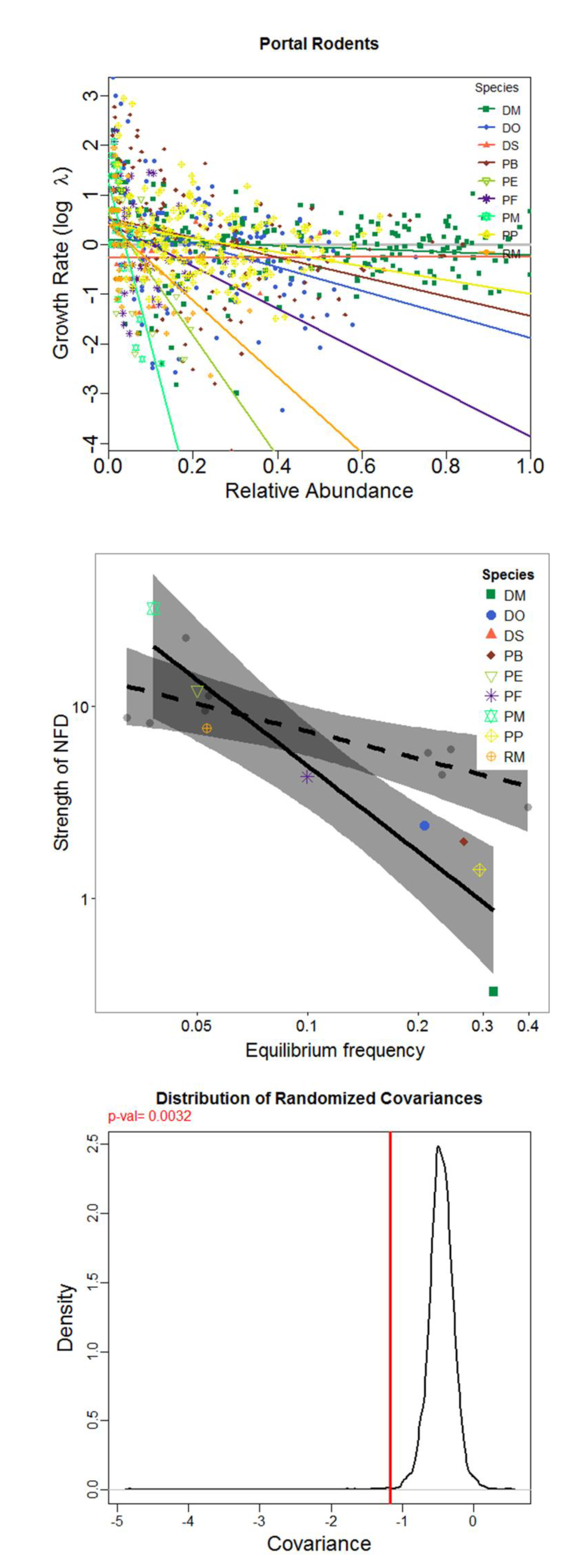
An example of our analyses using the Portal rodent community. (A) The relationship between relative abundance and growth rate is fit for each species. The slope of this relationship is the “Strength of NFD” for each species, and the x-intercept is the “Equilibrium Frequency.” (B) Using each species as a datapoint, the covariance between these two parameters (-1.17 in this example) is the value of interest, but it contains known biases. Randomizations of the time series were used to create a baseline expectation for each unique community, and the observed relationship was considered relative to this baseline (the Randomized Covariance, −0.46 in this example). Thus, we use the proportional difference in the empirical covariance from random (observed covariance/mean randomized covariance, −1.17/-0.46 = 2.54). The empirical pattern in this community (solid black line, with shaded 95% confidence intervals) compared to one realization from the randomized results (out of 5000 randomizations), accounts for the bias in the negative frequency dependence estimates introduced by measurement error and the artificial relationship this creates (dashed line, with shaded 95% confidence intervals). (C) P-values were calculated as the proportion of randomized values larger than the observed asymmetry value (-1.17, p-value = 0.0032 in this example).

## Dealing with Uncertainty in Abundance Estimates

Time series data with significant observation uncertainty can show the signature of negative density or frequency dependence even when none exists (Freckleton *et al.*, 2006; Knape & de Valpine, 2012) because the calculation can create a negative bias in the resulting population growth estimates. We designed a randomization procedure to determine which communities showed a significant negative covariance between equilibrium frequency and the strength of negative frequency dependence. We reshuffled abundances from the original community time series 5000 times for each species independently, and then repeated all estimation methods with the time-randomized data. This procedure maintains relative abundances and observed variability, but the frequency dependence detected in the randomized data is due to uncertainty alone. We compared the covariance estimated from the original data to the distribution of covariance estimated from the randomized data sets to estimate effect size and p-values. We report the difference in the observed pattern from the mean randomized pattern and calculate the p-value as the proportion of randomized pattern values that are less than or equal to the observed pattern. Simulations confirm that this approach accounts for known biases resulting from uncertainty in the observations, as well as sampling biases for rare species (see Supplementary Information for details). In addition to removing uncertainty bias, this has the added benefit of removing any effects of community structure (e.g. richness) that may create a bias in the pattern of interest, because it creates a randomized expectation for each community. This is a conservative approach, but it enables us to identify communities in which the observed relationship between rarity and NDF is extremely unlikely to arise due to chance or statistical artifacts. The significance of the asymmetric negative frequency dependence pattern was assessed using Strimmer’s approach (Strimmer, 2008) to estimate the number of true null hypotheses, controlling the false discovery rate at 0.1 (Benjamini & Hochberg, 1995, 2000; Verhoeven *et al.*, 2005). All analyses were performed in the R platform ((R Development Core Team, 2012) see Supplementary Information).

## Immigration or Competition?

While Yenni et al (2012) predicted asymmetric frequency dependence emerging from competition, it is possible that other mechanisms could generate the same pattern. While, only competition has been explored theoretically, it is possible to imagine scenarios where other processes could generate or modify negative frequency dependence patterns. We explored one of the more likely processes to generate asymmetric frequency dependence: immigration. Immigration is an important process influencing the persistence of species (Brown & Kodric-Brown, 1977) and is expected to have particular influence on populations of rare species, thus creating the potential for immigration to influence or even create patterns of asymmetric frequency dependence. We used both our NFD simulation and a more traditional Lotka-Volterra based model (Chesson, 2000) to explore how immigration influenced negative frequency dependence in common and rare species (Appendix S4). Results from both models were similar: immigration did not create patterns of asymmetric frequency dependence and, if anything, significant amounts of immigration decreases the ability of our method to detect true *in situ* asymmetric negative frequency dependence (models and results in Appendix S4).

## Asymmetric Patterns of Negative Frequency Dependence Appear to be Common

When compared to the randomized time series, 46% (41 of 90) of communities demonstrated a significant relationship between relative abundance and negative frequency dependence, indicating that rare species experience stronger negative frequency dependence than common species (Fig 3A, Table 1). While negative relationships between rarity and the strength of negative frequency dependence often arose in randomized data as well (Fig 2B, 2C), in the communities where the observed pattern was statistically significant, the stronger negative frequency dependence of rare species could not be explained solely as a result of bias related to uncertainty in the abundance estimates. These results indicate real differences between rare and common species in the sensitivity of population growth rates to the abundance of heterospecifics and conspecifics.

**Table 1:**
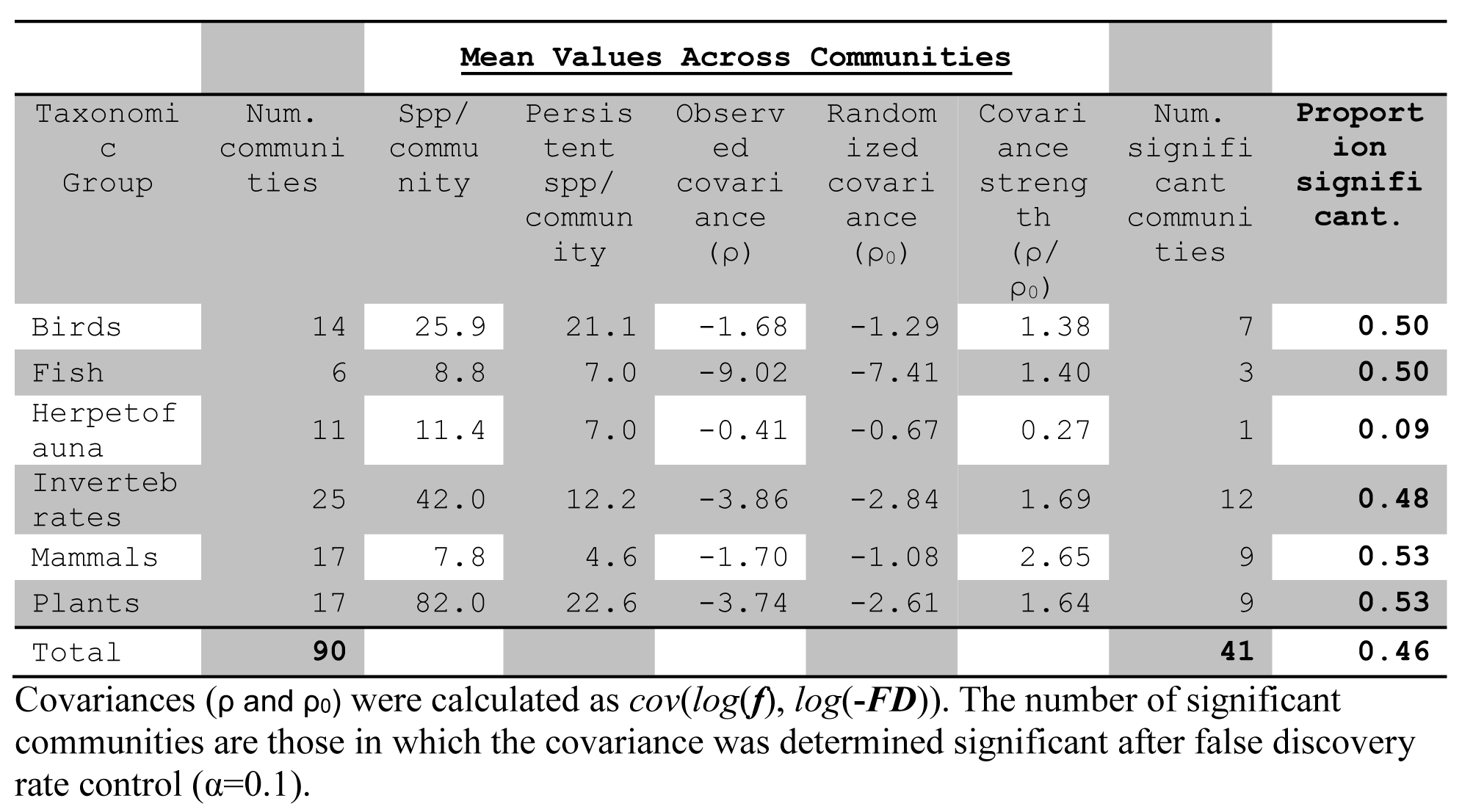
Summary results of community-level analyses by taxonomic group.

The prevalence of asymmetric negative frequency dependence within communities varied by taxonomic group (Fig 3B-G). In approximately half of bird, fish, invertebrate, mammal and plant communities (50%, 50%, 48%, 53% and 53%, respectively) rare species experienced stronger negative frequency dependence than their dominant counterparts (Table 1, Fig 3). However, only 9% of herpetofauna communities had a significant asymmetric negative frequency dependence structure. Why asymmetric negative frequency dependence is less prevalent in herpetofauna communities is unclear, though the documented sensitivity of herpetofauna to disturbance is one possibility for their differing dynamics (Collins & Storfer, 2003). Across all taxa, only a few communities had dominant species with stronger negative frequency dependence estimates than the rare species, but this pattern was never significant (see Table S1 in Supplementary Information). Thus, while rare species often, but not always, showed significantly stronger negative frequency dependence than common species, common species never showed stronger negative frequency dependence than rare species. Despite differences in diversity, complexity of species interactions and how species may achieve the necessary population dynamics, the phenomenon of rare species exhibiting stronger negative frequency dependence than more common species appears to be widespread across community types.

**Figure 3:**
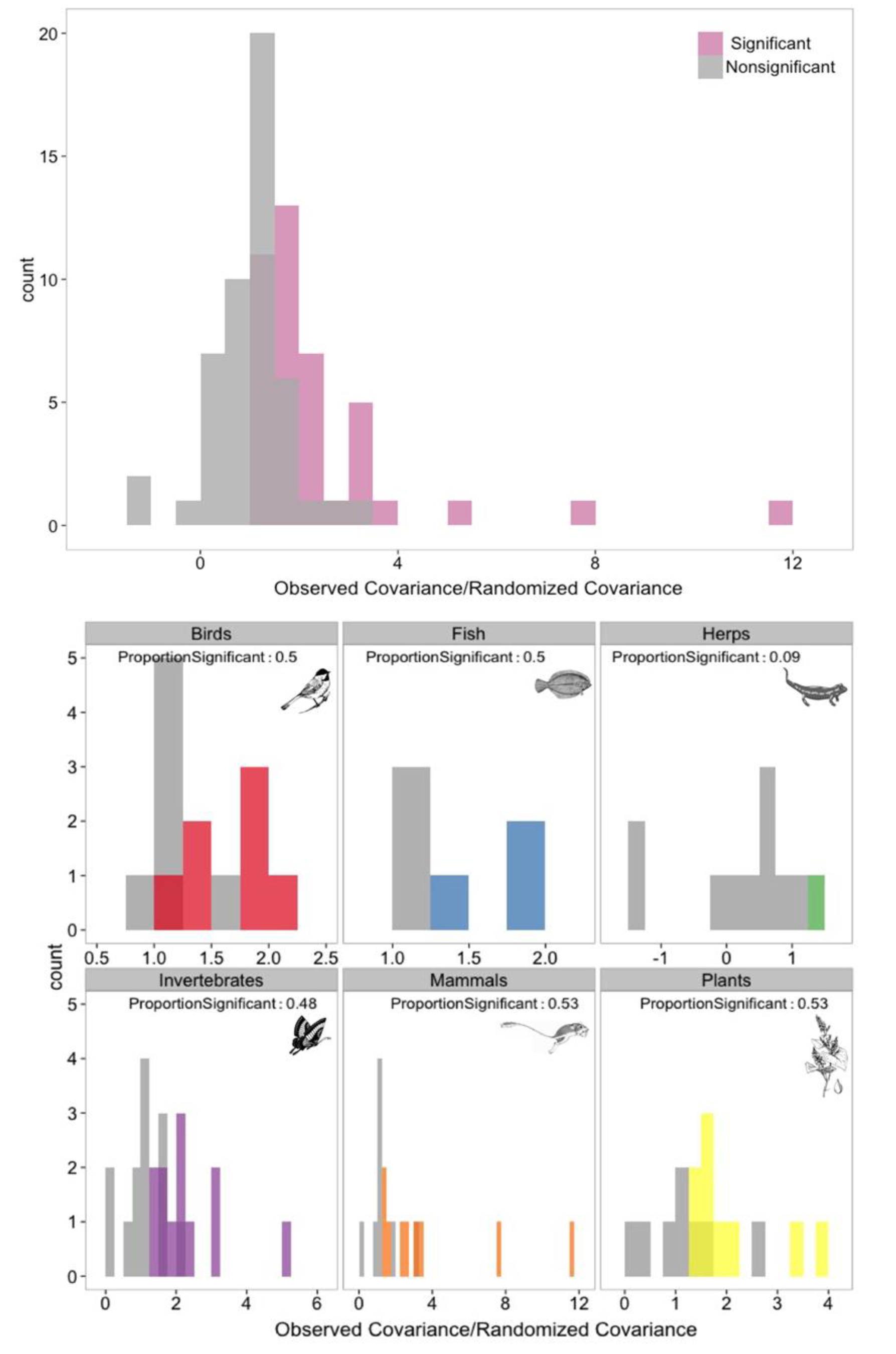
Results were compared to randomized data to account for bias introduced by measurement error and determine which relationships were significant (colored bars) or not (grey bars). P-values were calculated as the proportion of randomized values larger than the observed asymmetry value. (A) All results for the 90 communities included in the analysis. We detected a significantly asymmetric negative frequency dependence pattern in 46% of these communities. (B) Results separated by taxonomic group, showing the relative size of the asymmetry in negative frequency dependence between species in each community (Observed Covariance/Randomized Covariance). The proportion value in each panel is the proportion of communities in that group with a significant asymmetric negative frequency dependence pattern.

Simulations testing our randomization procedure for addressing bias caused by measurement error (Supplementary Information) demonstrated that our method consistently assigns significance when the relationship between equilibrium frequency and negative frequency dependence is strong, but it typically fails to detect weak relationships. Therefore, our non-significant results likely include some communities where a relationship between relative abundance and negative frequency dependence exists, but is too weak for our method to detect. If so, then the prevalence of strong negative frequency dependence in rare species is likely more common than our results indicate. Overall, our results suggest that in many communities, persistent rare species may commonly exhibit stronger negative frequency dependence than the more common species in the community.

## Implications

The theory developed in Yenni et al. (2012) and the results presented here imply that differences between rare and common species in their sensitivity to intraspecific and interspecific competition may explain the existence of persistent rare species in many communities. This finding leads to a variety of interesting research questions. Why are rare species more sensitive to intraspecific than interspecific competition? One possibility is that both strong sensitivity to congeners and an ability to increase rapidly when the population falls below its equilibrium frequency are traits of rare species, e.g. a result of specialization on a rare resource or susceptibility to virulent species-specific natural enemies. An alternative explanation is that strong negative frequency dependence is not a fixed trait but is context-dependent. Some species might show strong negative frequency dependence when conditions cause them to be rare members of the community but weaker negative frequency dependence in communities when they are more dominant. The limited availability of community time series data does not allow us to directly link strong negative frequency dependence to species traits or to test whether some species may show weak negative frequency dependence when they are dominant but strong negative frequency dependence when they are rare.

Yenni et al (2012) described a theoretical framework in which the stronger negative frequency dependence of rare species emerges from intraspecific competition. Our empirical results are consistent with the theory and suggest that asymmetric density dependence may structure many communities. Our simulations, provided in the Supplementary Information, further support the idea that the prevalence of asymmetric negative frequency dependence is not merely an artifact of uncertainty due to sampling biases or immigration. That is not to say that immigration as an ecological process, along with other processes such as facilitation and interaction networks, cycles, chaotic dynamics, and nonlinear negative frequency dependences could not also create or modify patterns of negative frequency dependence. For example, while we explored a particular implementation of immigration, it is still possible that immigration might be able to generate asymmetric negative frequency dependence under specific circumstances. Clearly it is important for coexistence theory to assess whether these other processes can also generate or enhance community-level patterns of asymmetric frequency dependence, especially as it relates to the concept of persistent rare species.

Regardless of the underlying mechanisms, or whether strong negative frequency dependence is a fixed or plastic trait of rare species, our results suggest that exploring the causes and consequences of asymmetric negative frequency dependence may be important not only for coexistence theory, but also for designing management strategies for rare species. If intraspecific competition is the primary cause of rarity, then indirect effects of environmental change mediated by competitive interactions (Adler *et al.*, 2012) might be relatively weak for rare species compared to common species. Similarly, we might expect management actions intended to increase the abundance of a rare species by reducing the density of interspecific competitors to be unsuccessful. However, if asymmetric negative frequency dependence is being generated by other mechanisms such as immigration, then the implications for designing management strategies will be very different. Investigating the mechanisms that lead to asymmetric negative frequency dependence is an important step for understanding, and potentially managing, the many rare species in natural communities.

## Future Analyses

Our analysis also highlights analytical and theoretical limitations to our understanding of negative frequency dependence that need to be addressed for research into negative frequency dependence to progress further. Definitive statements about the absolute strength of negative frequency dependence in common and rare species would require unbiased estimates of maximum population growth rates and density dependence. In contrast, our randomization test focuses on the relative strength of negative frequency dependence in common vs. rare species. Direct, unbiased estimates of negative frequency dependence would remove the need for our conservative approach, and could reveal the role of asymmetric negative frequency dependence in additional communities. An additional challenge for future research is disentangling the effects of intra‐ and interspecific competition from other potential mechanisms of negative frequency dependence such as immigration or nonlinear dynamics. Addressing this challenge will also require more sophisticated methods for correcting negative frequency dependence for bias or, alternatively, individual-level data on vital rates and dispersal.

## Summary

Our work suggests that strong negative frequency dependence in persistent rare species could be a mechanism that helps offset stochastic extinction risk. If true, this concept has wide-ranging implications for our understanding of coexistence. However, in exploring this idea, we found severe limitations in the current state of data, statistics, and theory that prevented us from conducting robust empirical analyses or interpreting rigorously the mechanism behind the patterns we uncovered. Given the importance, and possible generality, of asymmetric frequency dependence, we hope that our work here will inspire others to help solve these challenges so we may all better understand the mechanisms that allow rare species to persist in nature.

## Acknowledgements

A review by Dr. Robert Holt significantly improved the quality of this manuscript. Much of the credit for the Supplementary Information on negative frequency dependence goes to Dr. Holt. We thank all the researchers who have made their data publicly available and have permitted its use in this manuscript. Further information about these datasets and dataset specific acknowledgements of personnel and funding sources can be found in Appendix S1. Thank you to Sarah Supp and the Mammal Community Database, Elita Baldridge, Ryan O’Donnell, and all the contributors to the Ecological Data Wiki for assistance in identifying available community datasets. G. Yenni was supported by the Ecology Center at Utah State University and an NSF grant (DEB-1100664). PBA was supported by NSF DEB-1054040.

## Biosketches

Glenda Yenni is a multidisciplinary ecologist interested in drawing connections between theory and observation in community ecology. Peter Adler is a plant ecologist interested in explaining population and community dynamics in space and time. S.K. Morgan Ernest is a macroecologist and community ecologist studying the temporal dynamics of community assembly.

